# Bursts from the past: Intrinsic properties link a network model to zebra finch song

**DOI:** 10.1101/2024.05.18.594825

**Authors:** Nelson D. Medina, Daniel Margoliash

## Abstract

Neuronal intrinsic excitability is a mechanism implicated in learning and memory that is distinct from synaptic plasticity. Prior work in songbirds established that intrinsic properties (IPs) of premotor basal-ganglia-projecting neurons (HVC_X_) relate to learned song. Here we find that temporal song structure is related to specific HVC_X_ IPs: HVC_X_ from birds who sang longer songs including longer invariant vocalizations (harmonic stacks) had IPs that reflected increased post-inhibitory rebound. This suggests a rebound excitation mechanism underlying the ability of HVC_X_ neurons to integrate over long periods of time throughout the song and represent sequence information. To explore this, we constructed a network model of realistic neurons showing how in-vivo HVC bursting properties link rebound excitation to network structure and behavior. These results demonstrate an explicit link between neuronal IPs and learned behavior. We propose that sequential behaviors exhibiting temporal regularity require IPs to be included in realistic network-level descriptions.

## Introduction

Intrinsic excitability is a broad term that refers to a neuron’s membrane properties (ion channel density and composition) that regulate its firing properties relative to the strength of its inputs. This central feature of signal processing in individual neurons gives rise to a large diversity of functional properties^1–3,45,50^. For example, neurons that participate in rhythmic activity express specific channels (HCN and T-type Ca^2+^) that open at relatively hyperpolarized potentials and, in conjunction with release from inhibition (a network property), give rise to spike rebound excitation^4–6^ which then drives oscillations. As well as the types of ion channels expressed, regulation of excitability by means of ion channel density affects the input-output relationship of neurons^3,7,8^. Plasticity of these intrinsic properties (IPs), as well as activity-dependent plasticity within dendrites^8–10^, are related to the computational properties of neurons, networks, and behavior^17–23,51,52^. For example, IPs are implicated in mechanistic explanations for neuronal computation in the OFF-sensitive properties of retinal ganglion cells^11–13^ and in the encoding of species-specific acoustic communication in crickets^14–16^. In both cases, post-inhibitory rebound depolarization is a critical component of their integrative neuronal properties. Such observations motivate a search for the rules that relate the organization of IPs to behavior.

In order to investigate how neuronal IPs contribute to learning, network structure, and behavior, we leveraged the well-established model system of vocal production in songbirds. Specifically, we focused on the learned stereotyped song of the male zebra finch. The premotor nucleus of the song system, HVC, has two major classes of projection neurons (PNs): those projecting to the motor nucleus RA (HVC_RA_) and those projecting to the basal ganglia (HVC_X_) (Fig. 1A). A subset of these neurons is active when the bird sings^24–26^, emitting stereotyped and precisely timed spike bursts at specific moments of the bird’s song^26–29^. One distinction between HVC_RA_ and HVC_X_ is that the former only fire one burst per motif (canonical sequence of song syllables) whereas roughly half of HVC_X_ emit multiple bursts per motif.

**Figure 1.**
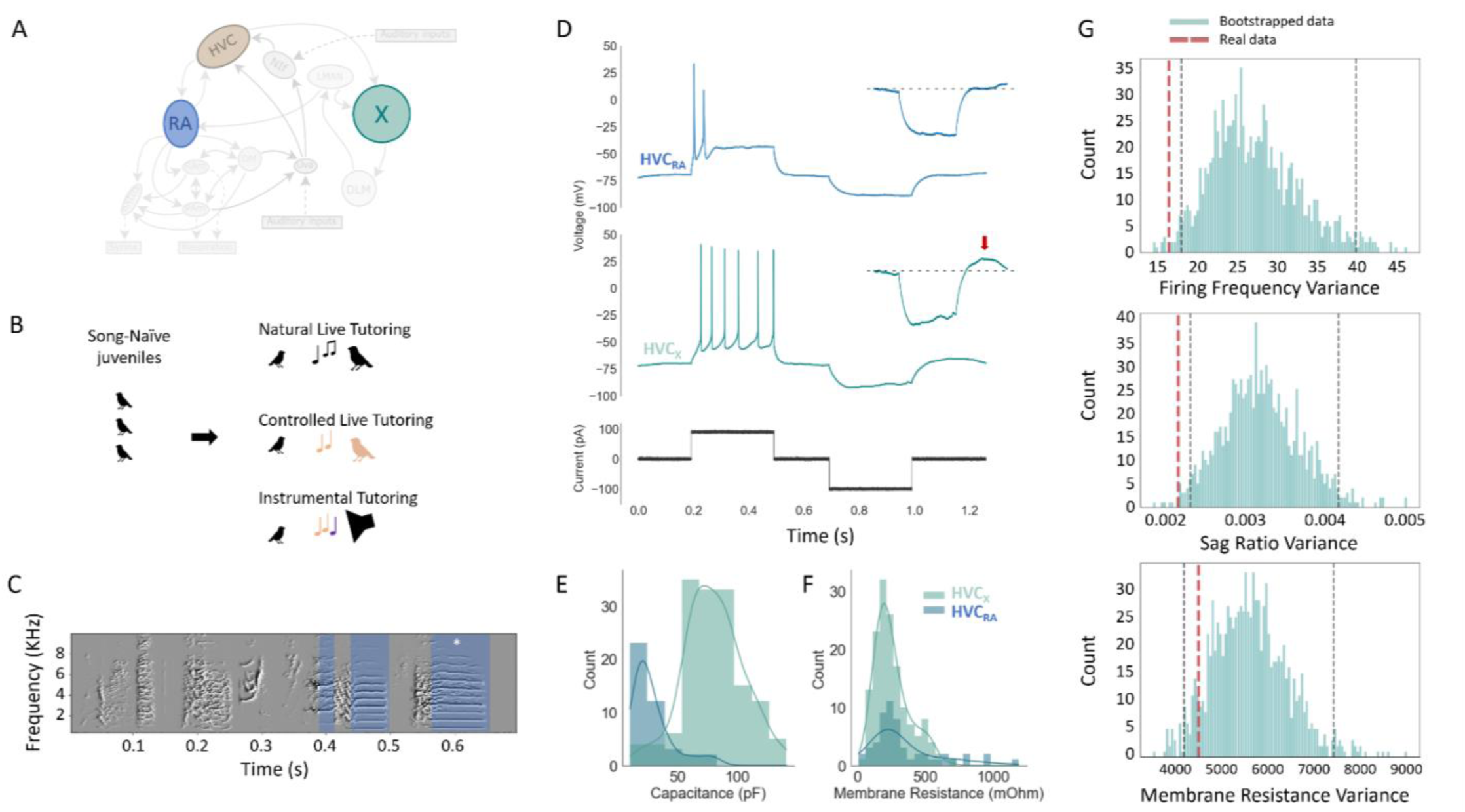
Intrinsic properties in HVC differ by projection classes and individual bird. **A**: Diagram of song circuit, illustrating two main projection targets of HVC (Area X and Nucleus RA). **B**: Illustration of our three tutoring paradigms. Natural live tutoring occurred in a family/sibling context; in controlled live tutoring juveniles interacted solely with a single adult male; in instrumental tutoring juveniles engaged a device (pulling a string) to elicit tutor song playback. **C:** An example motif sung by one bird, with harmonic stacks highlighted in blue and the longest stack marked by a white asterisk. **D**: Example traces from an HVC_RA_ (top panel in blue) and an HVC_X_ (middle panel in green), evoked by the current shown in black (bottom panel). Inserts show a zoomed in view of the hyperpolarized portion of the traces with a dotted line denoting the pre-inhibition baseline. For the HVC_X_ neuron, a post-inhibitory rebound response can be seen (red arrow). **E, F**: Histograms showing distributions of membrane capacitance and resistance, for 147 HVC_X_ (green), and 42 HVC_RA_ (blue). Solid lines show the kernel density estimation. **G**: Bootstrapped distributions for firing frequency, sag ratios, and membrane resistance for the mean within-bird variances (95% confidence intervals in black dashed line) compared to real mean within-bird variance (red dashed line).

Recent results demonstrate that the IPs of HVC PNs are developmentally regulated, depend on exposure to tutor song during the critical period for song learning, and vary by neuron class^18,30,31^. A detailed study of HVC_X_ IPs demonstrated that they are similar within individual birds and vary among birds in a way that reflects their learned song^31^. Birds who sing similar songs also express similar IPs. Disrupting auditory feedback during singing resulted in degraded fidelity of within-bird homogeneity of IPs. Thus, the configuration of HVC_X_ IPs is influenced by activity-dependent processes during development and in the context of adult song maintenance can be viewed as representing an error signal. However, whether specific features of IPs are related to specific spectral and/or temporal differences of individuals’ songs remains unknown. Here, we explore how IPs might interact with network architecture to encode complex learned behavior.

We evaluated the hypothesis that specific IPs are related to specific song features by manipulating the learning experience and found an explicit link between rebound excitability and temporal structure of song. We then used our experimental results to constrain a network model of Hodgkin-Huxley neurons that captures network properties of HVC_X_ neurons in the framework of spike rebound excitation. This network architecture provides a mechanistic framework with which to interpret previous work relating to syllable sequence selectivity in HVC and addresses timing delays in the VTA-basal ganglia projection that reflect song errors^32^.

## Results

In vitro whole-cell recordings were performed in 195 neurons from 38 adult male birds and across 3 tutoring paradigms (Fig. 1B) to assess neuronal intrinsic excitability and passive membrane properties. Songs were recorded once birds were older than 120 days, at which point one representative motif was chosen for analysis (Fig. 1C). HVC was identified as a dark myelinated region under brightfield illumination and confirmed with retrograde labeling using GFP or rhodamine (n = 4 animals, 8 slices). This definition of HVC was independently verified by the presence of canonical firing patterns of HVC projection neurons. HVC_X_ and HVC_RA_ were distinguished by their characteristic firing properties, as previously reported ^18,30,31,33–35^. HVC_RA_ neurons (n = 42) showed robust spike adaptation to depolarizing currents (100 – 150 pA square pulses, 10 pA steps, 300 ms), firing few spikes at stimulus onset riding atop a large, depolarized plateau (>20 mV) (Fig. 1D). HVC_X_ neurons (n = 153) had firing properties that included continuous firing with smooth spike after-hyperpolarization, modest spike adaptation, voltage sag and post-inhibitory rebound depolarization to negative applied currents (Fig. 1D, red arrow). The membrane capacitance of the HVC_X_ neurons was larger than that of HVC_RA_ (KS test, p < 1^-15^) (Fig. 1E,F)^33^, whereas there were no differences in the membrane resistance between the two classes (KS test, p = 0.21). Only the HVC_X_ showed post-inhibitory rebound^30^.

We used three designs to tutor birds. In the first, natural live tutoring design, birds were raised by their parents in individual cages while in acoustic contact with birds from other cages (see Methods). Such conditions tend to maximize song copying accuracy and leads to pairs of male siblings with small variations in their songs. For this design we chose birds opportunistically without regard to the features of the fathers’ songs. This design resulted in 14 male birds (4 families) from which we obtained electrophysiological recordings. Comparing the songs of all pairs of siblings by means of a commonly used song similarity measure^36^ yielded an average similarity of 85 ± 10% (N = 15 pairs; one bird that was a father was excluded from comparisons). Thus, we had strong song copying but weak control over the features of the tutor songs. In a second, controlled live tutoring design, birds

were raised in sound isolation chambers with their parents until 15-20 days post hatch (DPH), at which point the father was removed. Once male juveniles were identified, they were each placed in a separate sound isolation chamber with an adult male tutor. This provided for efficiency, allowing us to choose tutors whose song structure was advantageous to the experimental goals without requiring the tutors to successfully breed. This approach yielded 18 experimental birds (11 families with an average of 1.6 ± 0.98 siblings per family) from which we obtained electrophysiological recordings. Over the course of these experiments, a relation between features of song timing and HVC_X_ IPs emerged (described below). This motivated adopting a third design which allowed us to directly control the tutor songs that the juveniles heard. In this third, instrumental tutoring design, female–raised males at 32-40 DPH were each transferred to a sound isolation chamber where pulling a string provided instrumental access to song playback^16,32^. Electrophysiological recordings were obtained from 6 birds (4 families, 2 pairs of siblings) in this design. Combining birds from all three designs yielded a dataset of 38 birds.

Once a bird reached adulthood (≥ 120 DPH), we made slice electrophysiological recordings from HVC_X_ and evaluated their IPs. A notable feature of HVC_X_ is their relative homogeneity of IPs within a bird^31^. To assess this in our recordings, we first evaluated the within-bird IP similarity by bootstrapping features of the raw data. A subset of features of the intracellular traces correlated with the series resistance of the recordings (including spike amplitude but not firing frequency and sag ratio) and were thus excluded from analyses (see Methods). Bootstrapping consisted of maintaining the numbers of neurons per bird as appeared in the real data, then randomly reassigning neurons to one of 38 surrogate birds, repeating this procedure to generate 1,000 shuffled datasets. Thus, we evaluated the average within-bird variance for the real and bootstrapped data for firing frequency, sag ratio, and membrane resistance. The real neurons were significantly less variable within birds for firing frequency (p = 0.002) and sag ratio (p = 0.012) but not for membrane resistance (p = 0.269) (Fig. 1G). To address whether the apparent within-bird similarity arose artifactually from features of the recordings, we also performed a bootstrap analysis on the holding current, which is a good indicator of break-in and recording quality. We found no significant difference between the real data and the bootstrapped data for holding current (p = 0.102), indicating that variation in basic features of the recording quality do not explain the variation observed among birds.

### HVC_X_ intrinsic properties are related to temporal features of the individual’s song

Post-inhibitory rebound firing is often associated with rhythmogenesis^37^, which helped to motivate our hypothesis that the rebound excitation of HVC_X_ neurons could be related to temporal structure of song. To characterize features of song timing, for each of the 38 songs we examined the number of syllables, the duration of the song motif, and periods of continuous spectrally invariant vocalizations called harmonic stacks (blue shaded region, Fig. 1C). Initially, we focused on the longest duration harmonic stack in each song because it is the longest period of constant vocal output. Harmonic stacks are associated with relatively static syringeal muscle activity and respiratory pressure^59,60^, which could potentially be reflected in longer periods of temporal neuronal integration in HVC. We hypothesized that temporal integration processes within HVC would be more readily apparent studying these long harmonic elements. The number of syllables in the songs varied from 2 to 7, while song duration varied from 258 ms to 1487 ms. Songs had between 1 and 5 harmonic stacks, with the duration of the longest harmonic stack (asterisk, Fig. 1C) varying over 5–fold across the 38 birds (40 ms to 200 ms) (Fig. 2A, green points). Longer songs tended to include greater numbers of syllables as well as longer syllables and longer harmonics. We found that the duration of the longest harmonic stack was correlated with the duration of the remainder of the motif (motif duration minus the longest stack duration) (Pearson’s R: R^2^ = 0.47, p = 3.13 x 10^-6^) (Fig. 2A). This represents previously unreported temporal structure in zebra finch song, suggesting that longer songs preferentially include long harmonic elements. In our birds, we find that while longer songs tend to include more syllables (linear regression: R^2^ = 0.55) (Fig. 2C), they also tend to include longer syllables (linear regression: R^2^ = 0.37) (Fig. 2B). Lastly, we further confirmed such a relationship in a dataset of 52 songs from other labs^38^ (Pearson’s R: R^2^ = 0.24, p = 0.0003) (Fig. 2A, yellow points). Thus, zebra finches with longer songs tend to have longer syllables, including longer harmonic stacks, a result that builds on observations that zebra finch songs have isochronous organization^42^. Furthermore, as previously reported^61,62^, there was a strong tendency for the longest harmonic stack (in our 38 birds, green dots Fig.2A) to appear near the end of the motif, with 78% of songs having their longest harmonic stack in the second half of the motif, and 55% of songs having the longest harmonic stack in the last third of the motif, and this is the result of developmental song learning^62^.

**Figure 2.**
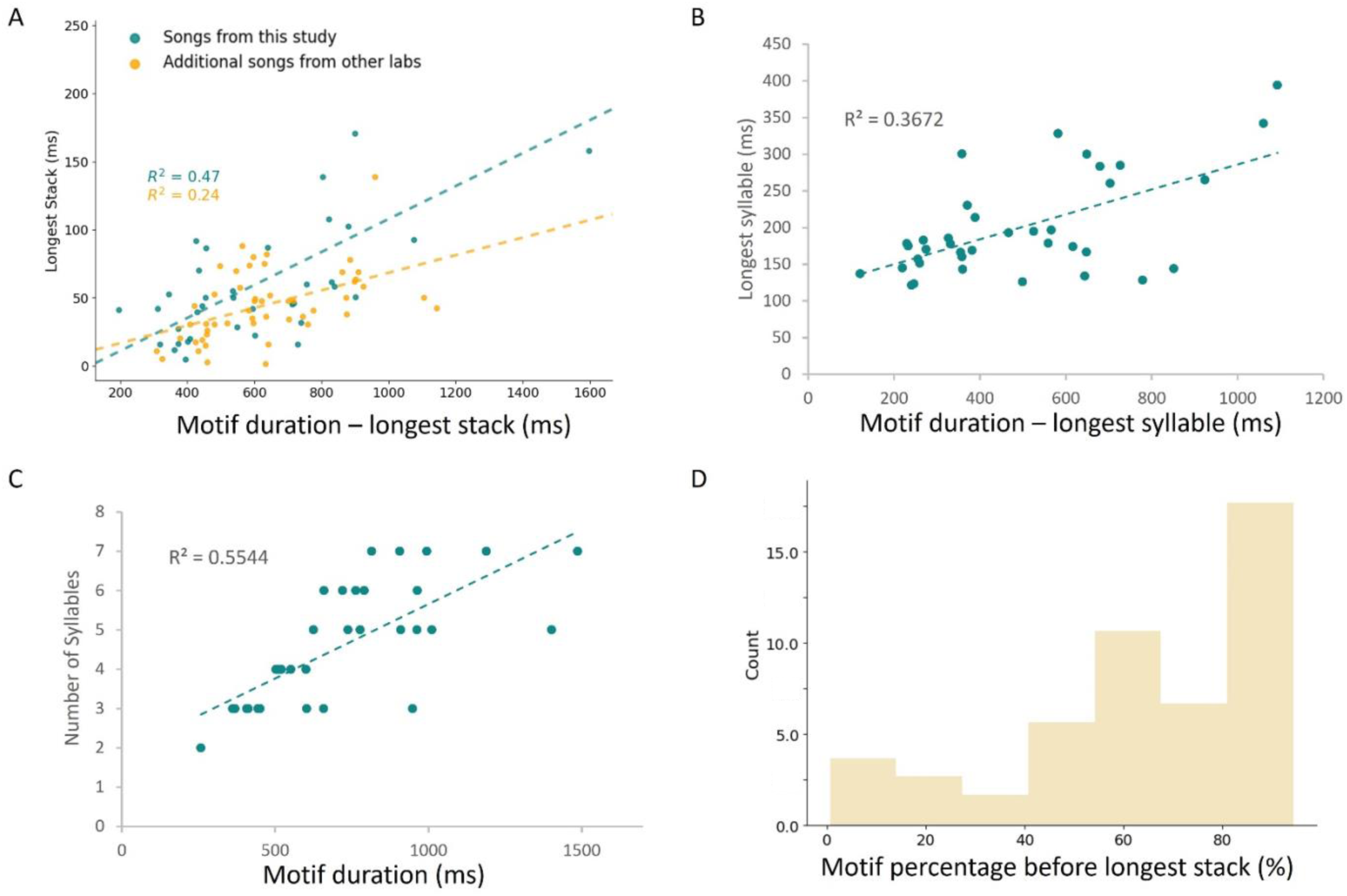
Internal temporal structure in zebra finch song. **A**: Durations of longest harmonic stacks in a song plotted against the remaining song motif (motif minus the longest harmonic stack) for songs in our study (green, R^2^: 0.47) and additional songs from other labs (yellow, R^2^: 0.24) and corresponding linear regression (dashed line **B, C:** Scatter plots for only the songs from this study (green) and lines of best fit for the number of syllables versus motif duration (R^2^: 0.55) and longest syllable versus motif remainder (motif minus longest syllable, R^2^: 0.37). **D:** Histogram (N=52 birds) showing the percentage of the song motif that occurs before the longest harmonic stack.

Having established these basic features of zebra finch song temporal structure, we explored the relationship between IPs and song timing by analyzing features of the recorded intracellular traces from birds who learned natural songs (i.e., non-modified songs from all three designs) (N = 33). We focused on the firing frequency (dotted line, Fig. 3A), hyperpolarized voltage sag (ratio of square and triangle, Fig. 3A), the post-inhibitory rebound area (dashed box, Fig. 3A), membrane capacitance, and resistance as our measures of IPs. The two measures of behavior were the duration of the longest harmonic stack (Fig. 3B) and of the motif duration.

**Figure 3.**
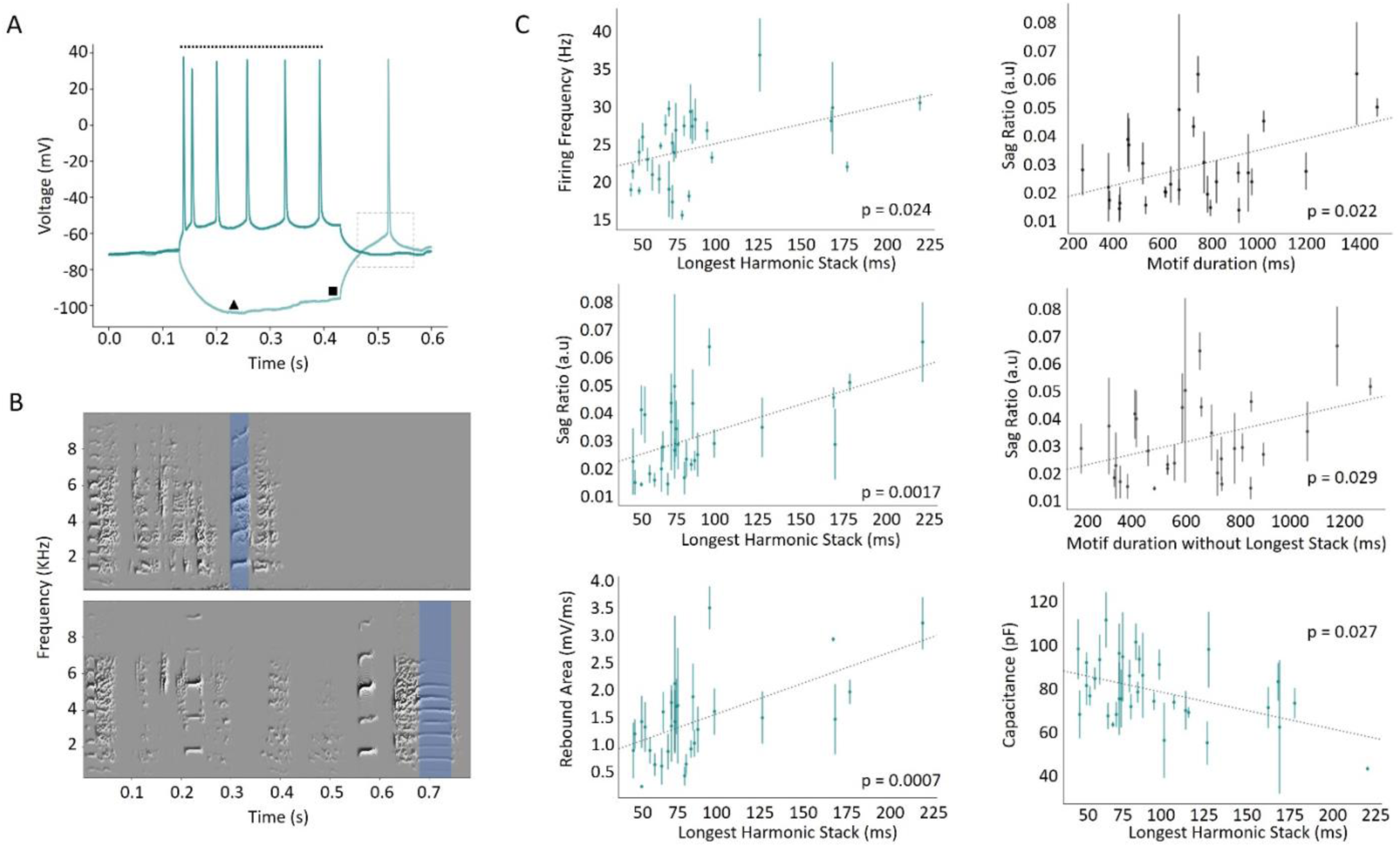
Intrinsic properties associated with rebound excitation are related to specific timing features of zebra finch song. **A**: Example voltage traces resulting from our standard current injections used to estimate cells’ post-inhibitory rebound excitability (black dotted box). Firing frequency was measured as the number of spikes over the duration of the spike train (dotted line), evoked by a 100pA depolarizing pulse (dark green, upper trace). Sag ratio was measured as the ratio between the minimum membrane potential (black triangle) evoked by a -100pA hyperpolarizing pulse (light green, lower trace), and the membrane potential before the release from the hyperpolarizing current injection (black square). **B**: Two example spectrograms showing representative song motifs of two birds. Blue shaded regions denote the longest duration harmonic stack in each song. **C**: Scatter plots of mean analyzed parameters for all HVC_X_ for each bird, against features of song duration (error bars are standard error of the mean); see text.

The firing frequency and mean sag ratio of HVC_X_ from each bird was positively correlated with the duration of their longest harmonic stack (Pearson’s R: R^2^ = 0.2, p = 0.024, and R^2^ = 0.32, p = 0.0017, respectively, Fig. 3C, left column, top and middle panels). We found that the average post-inhibitory rebound area correlated with the duration of the longest harmonic stack (Pearson’s R: R^2^ = 0.36, p = 0.0007, Fig. 3C, left column, bottom panel). The average sag ratio for each bird was also correlated with the motif duration (Pearson’s R: R^2^ = 0.2, p = 0.022, Fig. 3C, right column, top panel) and the remainder of the motif (motif duration – longest stack) (Pearson’s R: R^2^ = 0.17, p = 0.029, Fig. 3C, right column, middle panel). Lastly, average HVC_X_ membrane capacitance was inversely correlated (Pearson’s R: R^2^ = 0.22, p = 0.0027, Fig. 3C, right column, bottom panel), and membrane resistance was positively correlated (R^2^ = 0.2, p = 0.005, data not shown), with the duration of the longest harmonic stack of each bird. To further evaluate this result, we tested if the relationships between timing features and IPs we found were mostly driven by a few birds with longer duration longest harmonic stacks. Hence, we analyzed the data excluding birds who sang songs with longest harmonic stacks greater than 150 ms (Supplemental Fig. 2). We found that while some of the p values increased above 0.05 (p = 0.058 for rebound area vs. longest harmonic stack and p = 0.082 for sag ratio and longest harmonic stack), it remained significant for firing frequency and longest stack (Pearson’s R, p = 0.0017) and for sag ratio and motif duration (p = 0.024). However, when sag ratio was compared against the duration of the motif excluding the longest harmonic stack, there was no relationship (p = 0.85). Overall, this indicates similar structure in the core of the data set and in the birds with the longest harmonic stacks, with those birds accentuating several of the trends. Given that the birds with longest harmonic stacks also tended to have the longest songs, it may not be surprising that they had an outsized influence on some of these correlation measures.

Harmonic stacks are characterized by two principal features: by their duration in the time domain and fundamental frequency in the spectral domain. Hence, we also explored the relation of fundamental frequency with HVC_X_ IPs. In the 33 birds, the mean fundamental frequency of the longest harmonic stack was 779 ± 394 Hz. Overall, there was a systematic change in the direction of the best-fit lines when analyzing for fundamental frequency (cf. Fig. 3 and Supplemental Fig. 1), indicating some relation between fundamental frequency and IPs. There was, however, more noise in the correlations of IP measures with fundamental frequency as compared with correlations with time-domain measure. Most of the measures of IP failed to reach statistically significant correlations with the fundamental frequency of the longest harmonic stack (significance values ≥ 0.107), while firing frequency tended towards significance (R^2^ = 0.13, p = 0.63) (Supplemental Fig. 1). Higher firing frequency might be related to muscular effort hence tension of the syringeal membranes and fundamental frequency. Overall, these analyses supports our hypothesis that timing specifically is a central feature that is reliably represented by HVC_X_ IPs.

To explore how these results generalize to other temporal features of song, we also assessed IP relationships with the longest syllable duration and the sum of all harmonic stack elements. The longest harmonic stack was also the longest syllable in 11 of the 33 birds. Nonetheless, the longest syllable duration only tended towards significance when correlated with the average sag ratio for each bird (Pearson’s R: R^2^ = 0.12, p = 0.065, Supplemental Fig. 3, top left panel), which contrasts with the strong correlation when evaluating sag ratio vs. longest harmonic stack (see Fig. 3). The longest syllable did correlate with the average rebound area (Pearson’s R: R^2^ = 0.18, p = 0.0255, Supplemental Fig. 3, top right panel). The rebound area also correlated with the summed duration of all harmonic stack elements in the song (Pearson’s R: R^2^ = 0.4, p = 0.0003, Supplementary Figure 3, bottom right panel). This is expected given the multiple dependencies of the duration of different song elements, especially given that many long syllables are harmonics and that the sum of harmonics stacks (total duration) and the longest harmonic stack are strongly covarying (Pearson’s R: R^2^ = 0.64, p = 4 × 10^-7^, Supplemental Fig. 3, bottom left panel).

Lastly, we tested the relationship between temporal features of song and rebound excitation in data from Daou & Margoliash, 2020. The data was in the form of modeled ion conductances (gNa, gSK, gK, gh, gCaT). We regressed the average gh/gSK (related to voltage sag and firing frequency) on the duration of the longest harmonic stack for the 18 birds in that study. This yielded a positive correlation but did not reach statistical significance (Pearson’s R: p = 0.08, Supplemental Fig. 7). This could be due to a smaller number of birds (18 birds compared to 38), but possibly also to the underestimation of the HCN conductance, which is temperature dependent^56^. (Recordings in this study were conducted at elevated temperatures.)

### Song learning modifies intrinsic properties

Having established relationships between HVC_X_ intrinsic properties and features of song, we then evaluated two groups of birds whose songs were advantageous for elucidating the relation between harmonic stack duration and rebound excitation. Some of these birds were live tutored by the same tutor (second design) (Fig. 1B), others were trained through the instrumental tutoring protocol^36^ (third design) (Fig. 4A). With this approach, we generated two groups of birds (N = 5 birds per group) who sang very similar songs, except the songs of one group (group B, Fig. 4B) included an additional long harmonic stack. The modified tutor song incorporated an additional harmonic stack of 200 ms (chosen from a bird within our dataset) and inserted as the final syllable while adjusting the preceding gap to maintain the rhythmicity of the song (see Methods). All birds in group B acquired a song that had a terminal harmonic stack greater than 100 ms. Both tutoring paradigms yielded good song copying. The average song similarity to the original song was 96.5% ± 3.2% (N = 5) for group A and 98.6% ± 0.7% (N = 5) for group B, excluding the added harmonic stack. Additionally, the average similarity between group A and group B songs (excluding the added harmonic stack) was also high (95.6% ± 3.4%, N = 25). Thus, the instrumental tutoring approach focused the main changes in song to the long terminal harmonic stack syllable.

**Figure 4.**
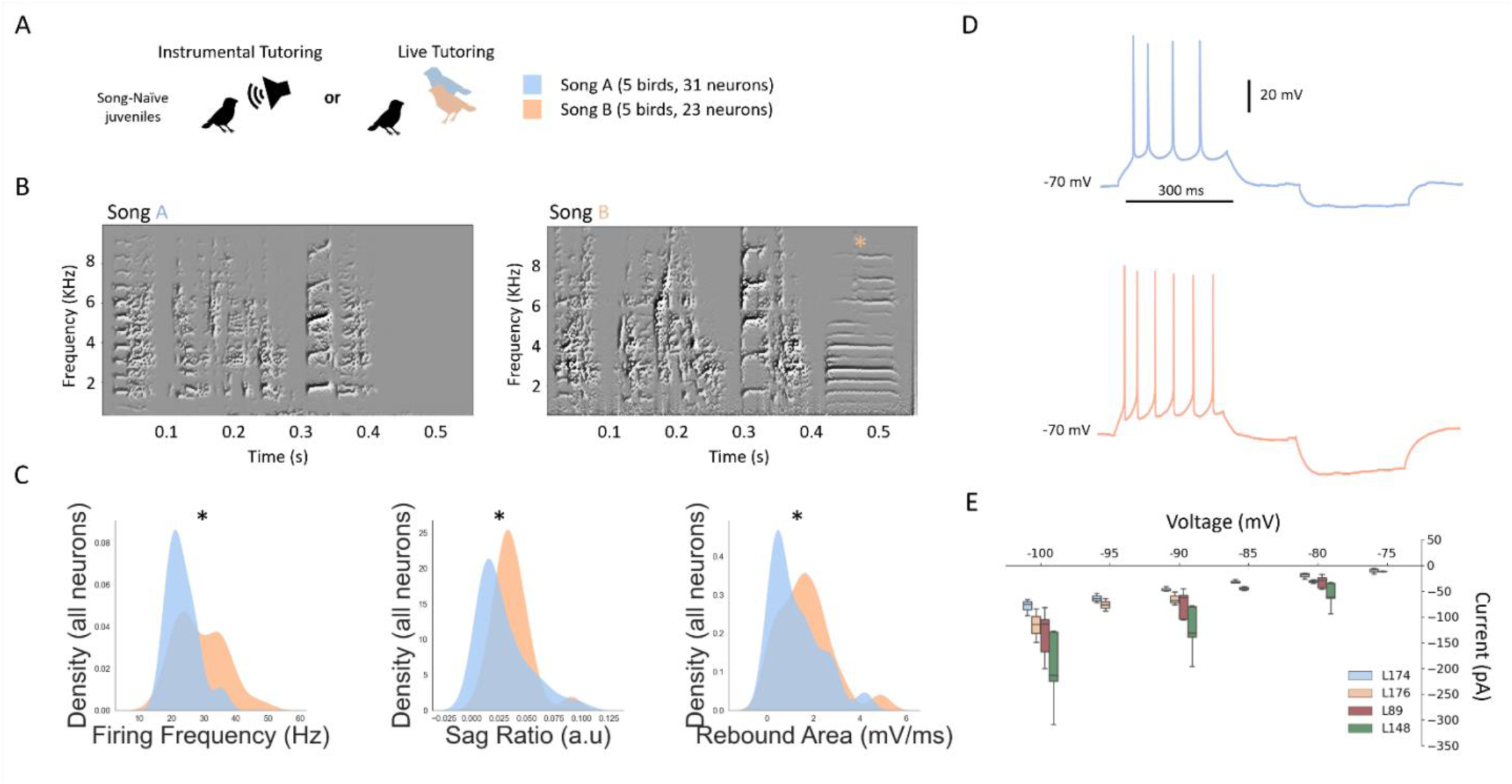
Instrumental manipulation of song learning changes intrinsic properties. **A**: Diagram of the instrumental and live tutoring paradigms that yielded birds singing very similar songs. Birds were tutored with either Song A or Song B. **B**: Example spectrograms from one bird from each song group. Song B included an added harmonic stack (orange asterisk). **C**: Distributions (kernel density estimations) for all neurons grouped by song type for evoked firing frequency, sag ratio, and membrane capacitance. Asterisks represent p < 0.05 for a KS test. **D:** An example voltage trace for a neuron from each song group (orange for modified song B, and blue for unmodified song A). Both neurons received the same input current of +100 pA for 300 ms followed by -100 pA for 300 ms. **E:** Average evoked currents for varying hyperpolarized voltage steps for four birds.

We then evaluated the IPs of HVC_X_ in the birds from the two groups. HVC_X_ neurons from birds who sang unmodified songs (N = 5 birds, 31 neurons), which had shorter harmonic stacks and shorter overall duration, had lower sag ratios (Mann-Whitney: p = 0.025), firing frequency (Mann-Whitney, p = 0.0051) and rebound area (Mann-Whitney: p = 0.0003) (Fig. 4C) when compared to those from birds who sang the modified longer song (N = 5 birds, 23 neurons). Lastly, we found that although the membrane capacitance trended towards smaller values for cells from the modified song group, it was not statistically significant (KS test, p = 0.08, data not shown). To address whether these differences were solely a product of changes in membrane resistance, we collected voltage-clamp data from four birds, two from the second experimental design, and two from the third design (N = 20 neurons). Two of the birds, L89 and L148 (longest stack durations of 91 ms and 66 ms, respectively), from the second live-tutored design, sang longer songs and had HVC_X_ with greater magnitudes of inward currents at hyperpolarized potentials (Fig. 4E, red and green bars). The other two birds, L174 and L176, came from the third design and sang songs A and B, respectively (Fig. 4B). L176’s average inward currents (Fig. 4E, orange bars) were consistently greater than L174’s (Fig. 4E, blue bars) across all membrane potentials below -70 mV. Notably, L148, who had the greatest evoked inward currents, had a total motif duration of 910 ms, while L89 had a motif duration of 738 ms.

Finally, we evaluated the contributions of heredity and song learning on IPs, using a mixed effects linear model (MLM). We examined the interactions between sag ratio, firing frequency, motif duration, the duration of the longest harmonic stack, and family as an indicator variable assuming random effect. Importantly, this also helped to assess which song features are the strongest predictors of IPs given that many of these covary. We ran two mixed effects linear models including birds from all designs (153 neurons, 4.1 ± 2.1 neurons per bird, 38 birds, 13 families), one using evoked firing frequency as the dependent variable, and another using sag ratio as the dependent variable. All values were normalized before running the model. Only the duration of the longest harmonic stack was significantly correlated with sag ratio (p = 0.018) and firing frequency (p = 0.008). To further investigate this, we analyzed sag ratio for a subset of families where all siblings were tutored with different songs. The average sag ratios of the siblings who learned different songs (13 birds, 4 families) were strongly correlated with the duration of the longest harmonic stack (linear regression: R^2^ = 0.6, p = 0.0002) but not the remainder of the motif (R^2^ = 0.1, p = 0.29, data not shown). Finally, we re-ran the model for sag ratio (still including family indicator and motif duration) but replaced longest harmonic stack duration with the total duration of harmonics in the song. In this case only the total duration of harmonics was significantly correlated with sag ratio (p = 0.021). Given that sum of harmonics and the longest harmonic duration are highly correlated (Supplemental Fig. 3, bottom left panel), we did not run a model including both.

In summary, we found that some temporal features of zebra finch song are correlated with IPs of HVC_X_ neurons, specifically, IPs that are associated with rebound excitability. We also found previously unreported internal structure in zebra finch song: harmonic stack duration, maximum syllable duration, syllable number and song duration are correlated. We also further described results showing that harmonic stacks dominate the endings of songs^59^. The totality of the results suggests that song temporal structure, and most reliably the duration of harmonic stack elements, most strongly predicts the differences among individual birds in HVC_X_ IPs. Our results do not exclude the possibility that spectral features of song, that also co-vary over the duration of song, are represented in the learned features of HVC_X_ IPs^62^.

### A network model of HVC leverages rebound excitation to reproduce HVC_X_ in-vivo properties

A major feature of HVC_X_ is their ability to integrate over a relatively long time (> 100 ms) while remaining sensitive to playback of sequences of song elements. For example, intracellular^39,40^ and extracellular^41^ recordings in anesthetized zebra finches have identified HVC neurons that encode multiple syllables. Numerous results are consistent with the hypothesis that these properties are also expressed during singing^24,26,53,57^ with detailed analyses of neuronal response properties limited to studies of sleeping or anesthetized birds.

Given our results linking features of song temporal structure to HVC_X_ rebound excitation, we hypothesized that hyperpolarization-activated current (Ih) interacting with rebound excitation is the mechanism that gives rise to both long integration and sequence selectivity. This is consistent with the earliest model for sequence sensitive HVC neurons which invoked release from inhibition^27^ and subsequent observations of hyperpolarization being predictive of the strength of syllable-sequence sensitive bursts of putative HVC_X_ neurons^39,40^. In this view, we conceptualized HVC_X_ as coincidence detectors of two events that define an interval (interval encoders). An HVC_X_ neuron encodes the occurrence of two events, first, a release from inhibition (releasing Ih as a depolarizing current), and second, an excitatory synaptic event. In this framework, each HVC_X_ has an integration window defined by the magnitude of rebound excitation, which we can visualize by looking at its rebound area during in vitro recordings (blue area under voltage trace, Fig. 5A). The resulting curve is a window in time when the two events can sum to produce an action potential (Fig. 5A, black trace in right panel). Neurons with different rebound excitability (varying expression of HCN, T-type calcium^2+^, and SK channels) can then have different delays, and narrow or wide integration windows for otherwise subthreshold events. We investigated this integration window in vitro in 6 neurons (4 birds) by varying the delay between release from inhibition and a small depolarizing input to generate a distribution of delays that produce action potentials (Fig. 5B, right panel). Two of these cells (Fig. 5B, left panel) illustrate how low and high rebound excitation can yield different delays and narrow or wide integration windows (blue and red traces respectively, Fig. 5B, right panel).

**Figure 5.**
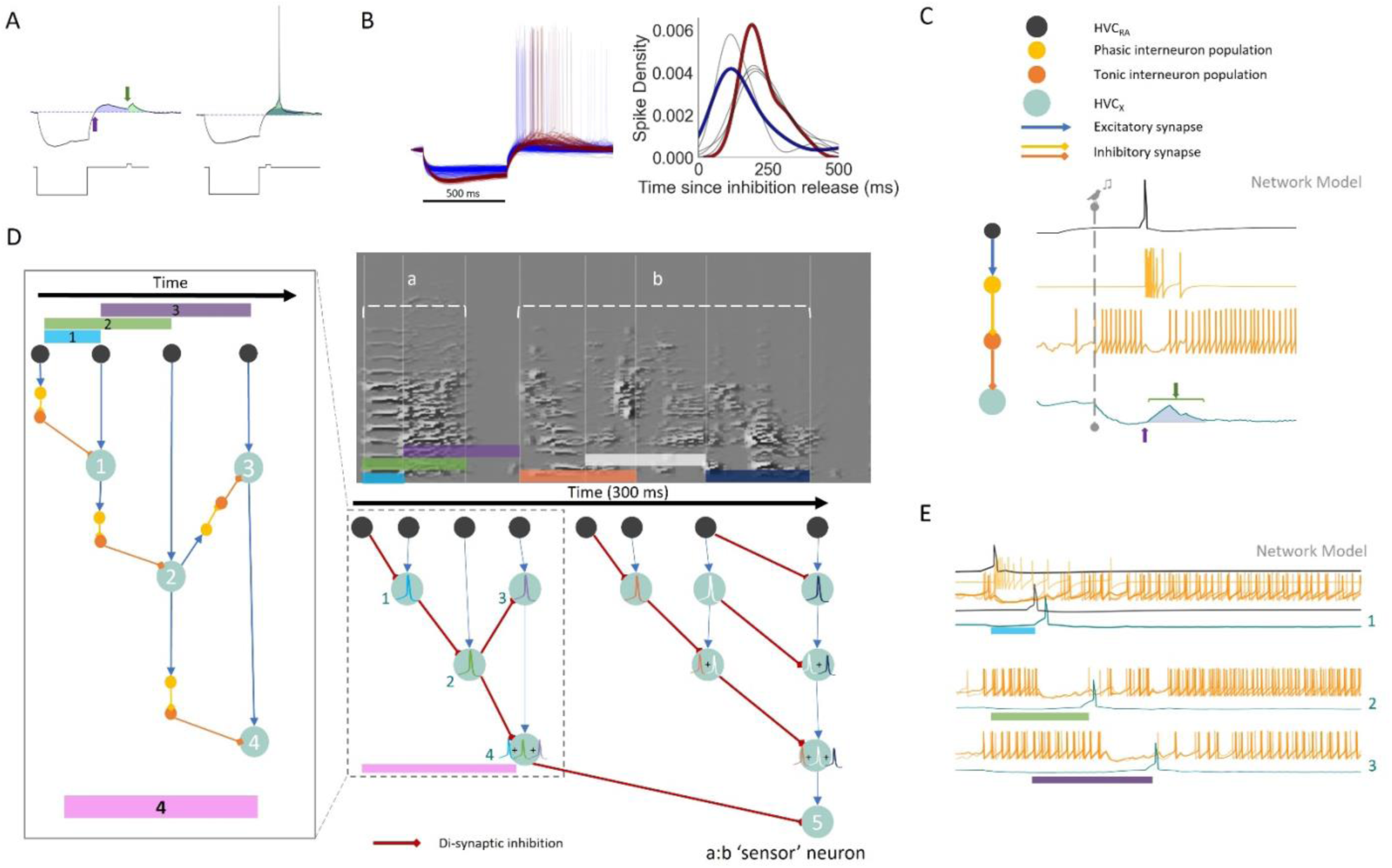
Hodgkin-Huxley network model leverages rebound excitation to produce in vivo bursting properties and sequence sensitivity. A: Experimentally recorded traces from neurons receiving hyperpolarizing and depolarizing currents at various delays (example current injection shown in bottom traces). The upper traces show voltage for rebound and direct depolarization timepoints (blue and green arrows, respectively) and highlights their area above resting potential (blue and green, respectively). Left panel shows example subthreshold responses, while right panel shows a suprathreshold response to a short delay (note overlap in blue and green areas). **B:** Traces from two HVC_X_ neurons with different sag and rebound responses to the same protocol in A. One neuron shows low magnitude rebound (blue) and another shows high magnitude rebound (red). Spike time distributions relative to delay for 6 neurons are shown on the right, including the example blue and red neurons shown on the left panel. **C:** Model diagram and Hodgkin-Huxley model traces for the basic sequence selectivity module (start of song depicted by dashed line with bird icon). The module utilizes inhibition release (blue arrow, bottom trace) and rebound, resulting in a depolarization ’window’ (blue highlighted area, bottom trace), and requires a second, depolarizing event during the rebound window (green bracket and arrow) to produce a spike (no spike shown). An HVC_RA_ neuron (top, black trace) excites a phasic interneuron (yellow trace), which inhibits a tonic interneuron (orange trace), resulting in disinhibition of the HVC_X_ neuron (bottom, green trace). Blue arrows represent excitatory synapses, yellow and orange arrows represent inhibitory synapses. **D:** Using the basic module from C, a backbone sequence of HVC_RA_ neurons define timepoints (vertical lines in spectrogram) in a two-syllable song segment. The timepoints define intervals that are encoded by spikes in HVC_X_ (green circles) which result from precise timing between disinhibition and excitation. The left inset shows a detailed view of the di-synaptic inhibition (red lines) in a portion of the greater circuit covering the entire two-syllable segment (small, dashed box). Spike waveform colors inside HVC_X_ circles correspond to colored intervals in song. **E:** Multiple modeled traces from neurons in this network, participating in interval representation for the corresponding colored intervals and numbered neurons in D.

We then constructed a network with cells modeled as Hodgkin-Huxley neurons with intrinsic properties representative of major HVC cell classes (HVC_RA_, HVC_int_, HVC_X_)^30^. We used the HH model previously reported in Daou & Margoliash 2020^31^, with additional differential equations for excitatory and inhibitory synapses (AMPA, NMDA, GABA). The model included the different sub-threshold changes that occur during singing for HVC_RA_ and HVC_X_, which are depolarized and hyperpolarized, respectively^34^. This was achieved by connecting HVC_X_ downstream of a population of GABAergic interneurons (tonic interneuron population Fig. 5C) whose activity increases at song onset. The increased background firing of the tonic interneurons was ‘hard-coded’ as an increased square depolarizing current that spanned the modeled motif duration. Similarly, an increase in input current was given to HVC_RA_ to simulate the depolarization seen at song onset. This defines a base circuit that reproduces static features of the network environment during singing. Next, we wired a di-synaptic (phasic) inhibitory connection^58^ between an RA projector and the tonic interneuron synapsing onto an HVC_X_ (Fig. 5C). This configuration depicts the simplest form of a module, without the secondary excitatory event to show the underlying curve that arises from inhibition release (blue area under HVC_X_ trace, Fig 5C). The duration and shape of the permissive window created by the gap in inhibition is determined in part by the magnitude of Ih and the return of inhibition. Building on this, a second RA projector that bursts later in time can then be wired with an excitatory synapse onto the same HVC_X_. This basic module produces a release from inhibition at one key timepoint (HVC_RA_ 1, Supplemental Fig. 4A) followed by excitation at another timepoint (HVC_RA_ 2, Supplemental Fig. 4A) in the HVC_X_, resulting in a spike only if HVC_RA_ 1 is followed by HVC_RA_ 2 at a certain delay (bottom panel, Supplemental Fig. 4A). Importantly, the disinhibition of the tonic interneuron by the phasic interneuron (Fig. 5C) can occur through presynaptic mechanisms and does not necessarily require direct somatic inhibition as shown here. Interneurons with different firing properties have been reported, and examples of tonic and phasic interneurons can readily be found in published work, such as Kosche et al. 2015^58^.

Thus, a module consists of one di-synaptic inhibitory connection and a monosynaptic excitatory connection onto the same HVC_X_. The monosynaptic excitatory input can originate from HVC_RA_ neurons, but also from other HVC_X_ neurons, as has been very recently described^63^. In the latter case, this results in a series of HVCX with nested dependencies that encode increasingly longer intervals with sequence specificity (Fig. 5D). The resulting network’s behavior includes a neuron which only spikes after the correct sequence of song associated HVC_RA_ bursts, with their appropriate timing (HVC_X_ 5 in Fig. 5D). In the model, the duration of gaps in tonic interneuron activity restricts the size of the intervals that can be encoded. Therefore, our model predicts that the gaps in tonic interneuron activity during singing or playback experiments should correlate with temporal song features, such as the duration of the longest harmonic stack or other song syllables/notes. This is consistent with results showing that gaps in inhibition are crucial for shaping neural sequences^58^. The model structure also predicts that there should be greater numbers of HVC_X_ integrating over shorter periods of time than HVC_X_ integrating over longer periods of time. To date, there is no such data available to test this prediction.

Here, we chose burst times for HVC_RA_ at biologically plausible syllable transitions^24,26^ that create intervals within a two-syllable segment of song and wired them in the modular fashion described above (Fig. 5D). In the model we describe, HVC_RA_ bursts form the backbone of the circuit and define key moments^24^; other models could incorporate key moments arising from other HVC_X_^26^. We placed HVC_RA_ bursts times at syllable onsets, offsets, and note transitions, as reported for subsets of HVC_RA_ neurons^24^ and HVC_X_ neurons^26^. While in principle burst times can be arbitrary in regard to the coincidence detection mechanism in our model and need not be anchored by song features, this would not be consistent with the sequence sensitive song system neurons described to date.

While roughly half of HVC_X_ burst only once, and are therefore captured by our model, the rest burst between two and four times (higher burst numbers are less common)^54^. In our framework, each burst of a multi-bursting HVC_X_ represents a separate detection of non-overlapping intervals. Thus, an HVC_X_ bursts for the detection of the first interval, returns to a hyperpolarized state, and then is able to burst again. Each burst is characterized by disinhibition (interval onset) followed by well-timed excitation (interval offset) (Supplemental Fig. 4B). Rapid sequential bursting of HVC_X_ would be a challenge to our model, but to date this has not been reported. Notably, because IPs determine the shape of the integration window within each neuron, multiple intervals encoded by the same neuron must be similar in duration. Hence, the modular structure of the network model allows for encoding multiple non-overlapping intervals by the same neuron. Finally, this configuration introduces delays associated with HVC_X_ bursting which align with delays in error signals observed during singing in inputs from the ventral tegmental area (VTA) onto Area X (see Discussion).

Lastly, we investigated the model’s flexibility in detecting small variations in timing by removing voltage-gated sodium channels from HVC_X_ and measuring depolarization without spiking. One HVC_X_ with inhibition release at a constant point (Supplemental Fig.5A, tonic interneuron gap) was tested against depolarizations from HVC_RA_ neurons at varying time points (Supplemental Fig. 5B, different colors represent different HVC_RA_ neurons). Under these conditions, the total size of the integration window for a single module (release from inhibition to return to baseline) is roughly 50 ms (Supplemental Fig. 5C, D).

## Discussion

During singing, at least some HVC projection neuron bursts are precisely time-locked to features of song^24,26^ and are sensitive to changes in timing. In anesthetized white-crowned sparrows, HVC neurons (likely HVC_X_, though it was not known at the time) require a sequence of notes before firing a burst of spikes at a transition point (e.g., the start of the next syllable)^27^. The ability for neurons to detect the sequence, represented by a burst of spikes, waned (fewer spikes) as the gap between syllables increased^27^. Thus, the strength of individual HVC_X_ spike bursts can reflect timing errors. The timing error refers to a difference in interval durations from some learned fixed point that is reliably predicted by song stereotypy. A similar experiment in zebra finches also showed sequence selectivity over hundreds of milliseconds^41^. Putative HVC_X_ neurons responded to sequential components of individual syllables, others required two syllables, while some only burst after presentation of long portions of BOS^39,40,41^.

Experiments in vivo have shed light onto the mechanisms underlying syllable selectivity. Sharp electrode recordings in zebra finches showed that selectivity is expressed in HVC_X_ neurons intracellularly as a hyperpolarization caused by playback of one song segment followed by a depolarization to a second segment^39,40^. Again, the projection identity of the neurons was not known at the time, but many of the neurons expressed hyperpolarizing potentials which are only expressed in HVC_X_^30^. During singing, different subthreshold mechanisms are involved in PN bursts^34^. HVC_RA_ neurons, that lack currents to support spike rebound bursting^30^, become depolarized when zebra finches sing, bringing their membrane voltage closer to their relatively high threshold. On the other hand, HVC_X_ become strongly hyperpolarized^34^, opening HCN channels^30^, which provide the inward current that drives post-inhibitory rebound. This dichotomy between PNs points to their functional differences and highlights an example of neuronal intrinsic properties giving rise to functional diversity in neural networks. More specifically, it points to rebound excitation being involved in HVC_X_ bursting, which requires specific intrinsic properties and network organization (timed inhibition onto the HVC_X_).

Our findings suggest that post-inhibitory rebound excitation in HVC_X_ could expand temporal integration. However, additional experiments are needed to directly quantify this effect in vivo. The relationship between IPs and behavior need to be evaluated in relation to network activity, which was not available to us for the cells we recorded. Yet, in other systems, such as in song recognition in crickets^14,15,16^ and OFF-ganglion cells in the retina^11,12,13^, rebound excitation serves as a mechanism that converts past inhibition into excitation with some delay. This mechanism enhances selectivity and precision in bursting. Specifically, it prevents early synaptic events from accidentally generating spikes while the neuron remains inhibited. As a result, the timing of integration for two events depends on the neuron’s intrinsic properties, the timing of inhibition release, and the gradual return of synaptic inhibition. The shape of the integration window allows small differences in timing to regulate the number of spikes in the burst. Consequently, the neuron’s bursts can provide information not only about the occurrence of the correct sequence but also about the relative timing of the encoded events. This is supported by the two–syllable experiments in white-crowned sparrows^27^ that showed HVC neurons’ burst strength reflected timing offset. These results show a surprising feature of IP involvement in network function, where not only is the presence of cell-type specific IPs important, but also that magnitude differences in IPs are reflective of network level computations and learned behavior. While previous results had shown individual variation in general IP composition, or experience-dependent ion channel expression, here we have linked rebound excitation directly to the temporal structure of a learned behavior.

The network model we constructed is able to replicate sequence sensitivity on the time scale of multiple syllables (Fig. 5) and, in principle, also retains the ability to detect small timing variations within individual coincidence detection modules (Fig. 5C). However, because the model is based on in vitro properties, it cannot perfectly reflect in vivo timing, and thus we limited model PNs to emit single spikes, not spike bursts. Furthermore, the model does not consider subcellular distributions of ion channels, which contribute to more nuanced responses in vivo^55^. Despite that, we found that the model remains sensitive to timing variations in the range of tens of milliseconds for a single module. Given that multiple modules can be nested together, this flexibility is extended by the number of HVC_X_ in a nested sequence. Further experiments are required to evaluate these properties under conditions of in vivo bursting and network effects. While this HH model provides a plausible framework for linking intrinsic properties to sequence propagation, it does not fully account for the observed relationship between IPs and song structure. A principal limitation constraining the current model is the absence of information for the same neurons combining characterization of both IPs and network activity during singing (or song playback), when HVC_X_ express activity related to song features. Addressing this gap would requires additional and challenging experiments and is beyond the scope of this study.

Nevertheless, if HVC_X_ neurons encode sequences as we propose, then the HVC to basal ganglia projection includes information about the relative timing of intervals, their order, and deviations from a learned stable point via the number of spikes in the burst. This supports the conclusion that HVC_X_ carry an error signal^31^. Furthermore, because HVC_X_ IPs show within-bird similarity, differences in EPSP timing downstream can be interpreted without added ambiguity from the presynaptic neurons’ excitability (a global solution). For example, a 50% variance in IPs (gh, gNa, gSK, gCa-T, gK) can generate large spike time differences (>100ms, Supplemental Fig. 5) in an HH model response to identical inputs. Thus, a global set of IPs promotes temporal precision and diminishes timing ambiguity at the population level. Different temporal features of song (duration of song elements and overall duration) would then give rise to different IPs in different birds. This provides an interpretation of previous results that showed song related clustering of HVC_X_ IPs in zebra finches^31^ and connects to findings about global temporal structure in zebra finch song^42^. This can explain why the duration of the longest harmonic stack was correlated with properties of rebound excitation, even though it is highly unlikely that the neurons we sampled were specifically associated with the longest harmonic stack during singing.

It is worth noting that the fundamental pieces of our model were previously reported by Lewicki & Konishi in 1995^39^ and 1996^40^, based on the cell circuit hypothesis by Margoliash in 1983^27^. However, the interpretation of those results lacked the greater context of the neuronal and network properties of HVC known today, specifically in the case of HVC_X_ neurons. Lewicki noted that, at the time, rebound excitation had not been observed in any cell in HVC during singing, or in vitro. Today, the rebound excitation of HVC_X_ neurons is well characterized and our results have connected it to temporal song structure and song learning. Their interpretation motivated a more complex model than present knowledge requires. Yet, Lewicki & Konishi were the first to demonstrate that inhibition caused by one element preceded a burst at the offset of the second element, and that the strength of the burst was predicted by the magnitude of the inhibition. Here we show how simpler modular synaptic structure uses rebound excitation to produce sequence selectivity.

At first glance our model may seem inconsistent with the recordings described in Lewicki & Konishi 1996^40^, where one syllable causes inhibition and another excitation, because our model assumes a ‘hard-coded’ constant inhibitory signal at song onset. This was motivated as a shorthand for the hyperpolarized in-vivo network environment of HVC_X_ during singing^34^. Instead, if every HVC_RA_ produces a burst in the tonic interneurons (orange neurons, Fig. 5) and inhibits downstream HVC_X_ (lateral inhibition), it can explain how isolated syllables produce strong hyperpolarization and provides a source for the song-induced hyperpolarization of the HVC_X_ population, which is currently not well-understood. Lateral inhibition between PNs has been previously described as common between HVC_RA_ and HVC_X_^34^, and monosynaptic excitatory and inhibitory connections between HVCX have only recently been reported^63^.

Finally, an important consideration about timing arises from our results. If each HVC_X_ neuron’s burst provides information about an interval only at the end of said interval, then information about errors that occurred early on in large sequences would be conveyed to the basal ganglia with a delay. Interestingly, VTA dopaminergic signals that arise from song errors arrive in the basal ganglia with a delay (∼60 ms)^32^. In our framework, each HVC_X_ burst signals to the basal ganglia that a song element was detected, providing the identity the element, and reflects information about whether a timing error occurred. This inherent delay aligns with the delay with which the VTA reports errors to the basal ganglia. A test of our interpretation is the prediction that the VTA delay should vary by bird and relate to the temporal structure of the song in alignment with such variation in HVC. This alignment in timing delays between HVC and VTA onto basal ganglia is an unintended consequence of our model architecture which, to the best of our knowledge, is the first attempt to address the significant delay between song errors and VTA-basal ganglia activity.

In conclusion, our study demonstrates the first explicit link between specific features of song and IPs of HVC PNs, potentially shedding light on the underlying mechanisms of both song production and song perception. Although harmonic stacks provide a useful test case for studying temporal integration, our findings suggest that IPs are broadly linked to song duration and structure, rather than specific syllable types. The modular structure of our network model offers new perspectives on how HVC_X_ encode sequences and highlights the importance of the synergy between intrinsic and synaptic neuronal properties in shaping network dynamics. Such mechanisms are likely to be more broadly represented than is currently understood.

## Acknowledgments

We thank Drs. Berthold Hedwig, Shivang Sullere, Ruth Anne Eatock, and Christian Hansel for their feedback on an earlier version of the manuscript and Dr. Arij Daou for sharing MATLAB code and general discussion on network modeling. Funding was provided by NIH 1UF1NS115821.

## Author Contributions

Conceptualization and design, both authors; supervision of research, D.M.; electrophysiology, N.M.; animal handling and training, N.M.; network modeling, N.M.; writing- original draft, N.M.; writing- conceptualization of structure, review, and editing, both authors.

## Methods

### Animals

All procedures were performed in accordance with regulations for animal testing and research and approved by the University of Chicago Institutional Animal Care and Use Committee (IACUC). Zebra finches (*Taeniopygia guttata*) were obtained from our breeding colony and housed on a 14:10 h light/dark cycle. Food and water were provided *ad libitum*. All birds were adults older than 120 days.

### Natural song learning

In this design, zebra finches were raised by their parents in individual cages where they could hear birds from other breeding cages as well as nearby flight aviaries. Birds were housed with their siblings and parents until 80 days post hatch (DPH), at which point they were moved to a flight aviary. Adult male zebra finches whose parentage was known were collected from our colony and used for experiments.

### Controlled song learning

Zebra finches were bred in sound attenuation chambers with their parents and siblings only. The father was removed when hatchlings reached 15-20 DPH, and house separately. At 40-45 DPH, juvenile males were identified by their chest and cheek plumage and separated to another sound attenuation chamber for song tutoring.

We took two approaches to control song tutoring. Live tutoring involved the introduction of an unrelated adult male into the sound chamber of a song-naïve juvenile. Tutor and tutee were housed together until the tutee reached 90 DPH. A second approach using instrumental tutoring occurred in the absence of other birds. We used triggered playback of pre-recorded song through a speaker, controlled by the software Sound Analysis Pro (SAP)^36^. Sound files used for instrumental song learning contained one three-motif bout of either naturally occurring song, or a manipulated song. In cases where manipulated songs included an added harmonic stack, we maintained the overall rhythm by using a previously developed method for finding rhythm in song^42^. In cases where multiple male siblings were old enough for tutoring, we placed them with different tutors.

### Sound recording and similarity analysis

Adult males were individually housed in sound attenuation chambers. Sound was collected with a microphone inside the chamber, amplified and digitized (Behringer UMC1820), and stored using SAP. Song similarity was accessed for pairs of single motifs using SAP’s mean value symmetric similarity after amplitude thresholding to capture all individual syllables.

A representative motif for each bird was chosen from song bouts that occurred in the days prior to slice experiments, and all analyses were performed on that representative motif. The representative motif was defined by the most complete version of the repeated syllables over many motifs (50-100). Distance calls between bouts and notes that only appear on the last motif in a bout were not included (potential calls). Hand labeling of motifs, syllables and gaps were verified in a semi-automated way using Chipper^46^ while harmonic stack durations were verified algorithmically with custom code that relied on Resin (Margoliash Lab Github), which identified moments of high amplitude and low frequency modulation (FM) by smoothing the time varying FM and placing an arbitrary threshold at 0.2 FM.

### Retrograde labeling

Birds were anesthetized with isoflurane and head-fixed using a stereotaxic frame. Bilateral craniotomies over Area X, or nucleus RA were made using predetermined coordinates (relative to Y Sinus with head angle of 18°) in mm: Area X: 1.5–2 lateral, 3.5 rostral, 3.5 deep, RA: 2.3–2.4 lateral, 1–1.1 caudal, 1.5–1.9 deep). Three birds were bilaterally injected with retrograde tracer (tetramethylrhodamine dextran from Invitrogen, 10-20 nL) using a Nanoject 2.0 or Nanoject 3.0. Similarly, in one bird we injected a self-complementary AAV carrying a GFP construct (sc-AAV9-GFP, UNC Vector core, 300-350 nL at 5nL/s) into Area X, and tetramethylrhodamine into RA. Slice experiments were conducted 7–10 days after injections to allow for transport or viral expression. Retrogradely labeled cells within HVC were identified with epifluorescence during recordings and used to confirm the identification of HVC using wide-field illumination.

### Slice preparation

Birds (n = 38) were anesthetized with isoflurane, checked with a foot pinch and then decapitated. A razorblade was used to block the brain through the skull. The blocked brain with skull still attached was placed in ice-cold sucrose-ACSF (225 mM sucrose, 2 MgSO_4_, 2 CaCl_2_, 1.25 NaH_2_PO_4_, 26 NaHCO_3_, and 10 glucose, 3 KCl, ph: 7.2–7.3) while the brain was removed. Horizontal slices (200–300 µM) were cut from both hemispheres using a vibratome (Series 1000, PELCO) and moved to incubate in a warm (32–34 °C) NMDG-ACSF (93 NMDG, 2.5 KCl, 1.2 NaH_2_PO_4_, 30 NaHCO_3_, 20 HEPES, 25 glucose, 5 sodium ascorbate, 2 thiourea, 3 sodium pyrubate, 10 MgSO_4_.7H_2_O, 0.5 CaCl_2_.2H_2_O) for 10-15 minutes before being moved to another incubation chamber containing standard recording ACSF (124 NaCl, 2.5 KCl, 1.2 NaH2PO_4_, 24 NaHCO_3_, 5 HEPES, 12.5 glucose, 2 MgSO_4_.7H_2_O, 2 CaCl_2_.2H_2_O). All ACSF solutions were bubbled continuously with 95% O_2_ 5% CO_2_. Slices were left to incubate for at least 30 minutes before being moved to the recording chamber.

### Whole-cell recordings

Recordings were done at moderately elevated temperatures (28-32 °C, controlled with an inline heater). Recordings were made using a Multiclamp 700B and digitized with a Digidata 1550B at 50 kHz (Axon Instruments), using a 10 kHz low-pass Bessel filter. HVC was first identified under wide-field illumination as a dark region in the horizontal slice at 4x magnification, and then by the presence of the characteristic electrophysiological responses of the two main classes of projection neurons in HVC (verified with retrograde labeling in 4 birds). Data collection was controlled using pClamp 10.4 (Molecular Devices). Recordings were made with fire polished glass pipettes (4-8 MΩ) and visually guided using a camera (Olympus Oly-150IR, or Hamamatsu Orca Fusion C15440). Electrodes were filled with 100 mM K-gluconate, 5mM MgCL_2_, 10mM EGTA, 2mM NA_2_-ATP, 0.3mM Na_3_-GTP, 40mM HEPES (pH: 7.2-7.3, osmolarity 290-300 mosM).

Whole cell recordings were performed in 195 HVC_X_ neurons, from 38 birds (mean neurons/bird = 5.13). A giga- Ohm seal was formed before break-in for all cells, and a period of 1-2 minutes was given before presentation of experimental protocols. Series resistance, membrane resistance, and time constant were calculated from a -10 mV step in voltage clamp. Series resistance was calculated as R_s_= V/I_peak_, membrane resistance was calculated as R_M_ = ΔV/I_steady state_, tau was calculated by fitting a polynomial (python polyfit) to the current trace and finding the time where the current reached 63% of its steady-state value. Capacitance was measured as CM = tau/RM. The median series resistance of cells was 22 MΩ ± 13.39 (73% of cells had R_S_ < 30 MΩ). Cells were then held to -70 mV in current clamp and presented with depolarizing and hyperpolarizing square current injections of varying amplitudes (-120 – -60 pA hyperpolarizing, 100 – 50 pA depolarizing, 300 ms duration). Firing frequency was calculated as the number of spikes divided by the time between the first spike and the last spike. Sag Ratio was calculated as the ratio between the most hyperpolarized voltage during the hyperpolarizing current injection and the voltage before release of hyperpolarization. Rebound area was calculated as the total area of the resulting rebound depolarization above pre-inhibition baseline for a 500 ms period after release from hyperpolarization. Sag ratio was calculated from the response to a hyperpolarizing -100 pA current injection as SagRatio = (Minimum voltage – Final Voltage)/Minimum voltage.

In a small number of cells, we injected a longer hyperpolarizing current step (500 ms) followed by a variable gap (10 – 390 ms, increasing by 20 ms), then injected with a 50ms depolarizing current step. The amplitudes of both current injections were adjusted on a per cell basis to keep them subthreshold for spiking when presented alone. In 20 cells from 4 birds (3,3,5, and 9 cells respectively), we performed voltage clamp experiments to produce I/V plots, stepping voltage from -100 mV to -60 mV, with step sizes of 10 or 5 mV.

All cells that fired consistent trains of spikes with spikes passing 0 mV were included in analysis. Features of cells that significantly varied with R_S_, assessed by linear regression, were not considered for analyses. Spike amplitude correlated with R_S_ (p < 0.004) and thus was excluded along with half-spike width, which depends on spike amplitude. Sag ratio and firing frequency did not correlate with R_S_ (p = 0.43 and 0.4, respectively).

### Hodgkin-Huxley Model

We used a single compartment conductance-based Hodgkin-Huxley model of HVC_X_ neurons to simulate the bursting properties of HVC neurons. The model includes the following currents: I_K_, I_NA_, L-type Ca^2+^ (I_ca-L_), T-Type Ca^2+^ (I_ca-T_), I_SK_, A-Type K^+^ (I_A_), and I_h_. HVC_RA_ and HVC_INT_ HH models were based on the HVC_X_ equations but with adjusted conductance values. HVC_RA_ for example had increased gSK values, and HVC_INT_ had decreased gSK values and increased gH. We updated the equations for the HCN channel (I_h_) to include voltage-gating behavior as reported in Boulet & Bruce^45^. Synaptic connections between neurons were modeled using the following formulations:

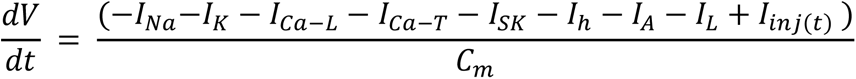

Equation 1. Hodgkin-Huxley single compartment neuron, adapted from Daou et al. 2013. Currents included: Voltage gated sodium (I_Na_), Voltage gated potassium (I_K_), L-type calcium channel (I_Ca-L_), low threshold activated T type calcium channel (I_Ca-T_), small conductance calcium dependent potassium channel (I_SK_), hyperpolarization activated cyclic nucleotide channel (I_h_), A-type potassium channel (I_A_), a leak current (I_L_) and any additional injected current provided, typically to model experimental current injections (I_inj_).

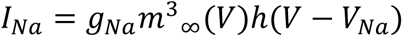

Equation 2. Example of ionic current equation for voltage-gated sodium channel. The equation includes terms for maximal channel conductance (g_Na_), activation parameter (m), neuron voltage (V), inactivation parameter (h), and the driving force as the neuron voltage (V) minus the reversal potential of the ion (V_Na_).

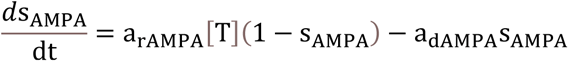

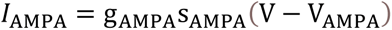

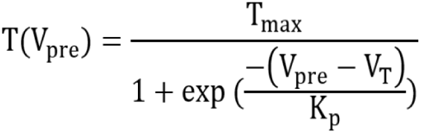

Equations 3-5. Example of the set of equations used to model ligand gated ion channels. Within the model, the equations used included AMPA, NMDA, and GABA_A_ and all depend on the pre-synaptic voltage. Tmax represents the total maximum transmission which is related to the presynaptic voltage (V_pre_) and the voltage for synaptic release (VT). The current provided by the receptor (I_AMPA_) depends on the maximal conductance (g_AMPA_) how much release there is at the synapse (S_AMPA_) and the driving force, modeled as the neuron’s voltage minus the reversal potential of the channel (V-V_AMPA_). The amount of synaptic transmission is determined by T.

Custom MATLAB code was used to run the network model. Rather than running the entire simulation at once for the large network (Fig. 5D), multiple smaller runs with adjusted timepoints were used, focusing on a single HVC_X_ neuron at a time. Each smaller run of the model contained 47 neurons (12 HVC_RA_, 11 delay neurons, 24 HVC_INT_, and 1 HVC_X_) and their synaptic connections for a total of 560 differential equations.

### Quantification and Statistical Analysis

Data files were exported from pClamp as axon binary files (.abf), which included multiple sweeps of a square wave current injection protocol (Fig. 1D, black trace, bottom panel). Files were accessed using a custom python library (pyABF, written by Scott W Harden: https://github.com/swharden/pyABF)^47^, and the raw traces were saved in Bark format (Margoliash Lab Github). Features of the raw data were extracted and analyzed using custom python code.

Linear mixed-effects models were implemented in two cases where we investigated the effect of song features and family (indicator variable) on IPs of 153 HVC_X_ neurons. For all linear regressions a Pearson’s R measure was used. Statistical comparisons of distributions were performed with the Kolmogorov-Smirnov test (KS test). All distributions are reported as mean ± standard deviation unless otherwise noted. Error bars in scatter plots are reported as the mean standard error.

## Supplemental Figures

**Supplemental Figure 1.**
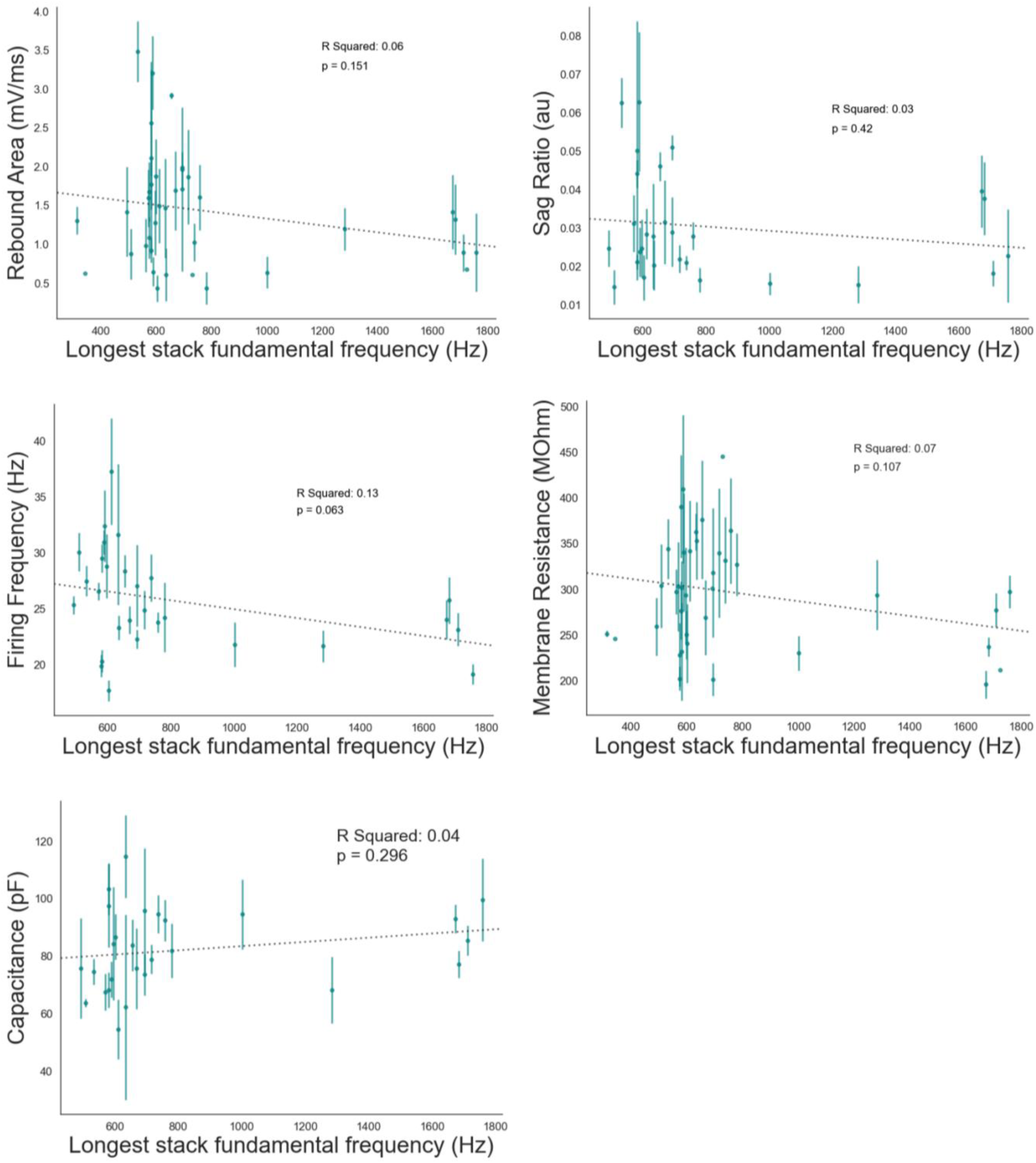
Intrinsic properties are unrelated to fundamental frequency of longest harmonic stack. Scatter plots of mean analyzed parameters for all HVC_X_ for birds singing natural songs, against the fundamental frequency of the longest harmonic stack. Each point is the mean value for each bird, and error bars represent standard error of the mean.

**Supplemental Figure 2.**
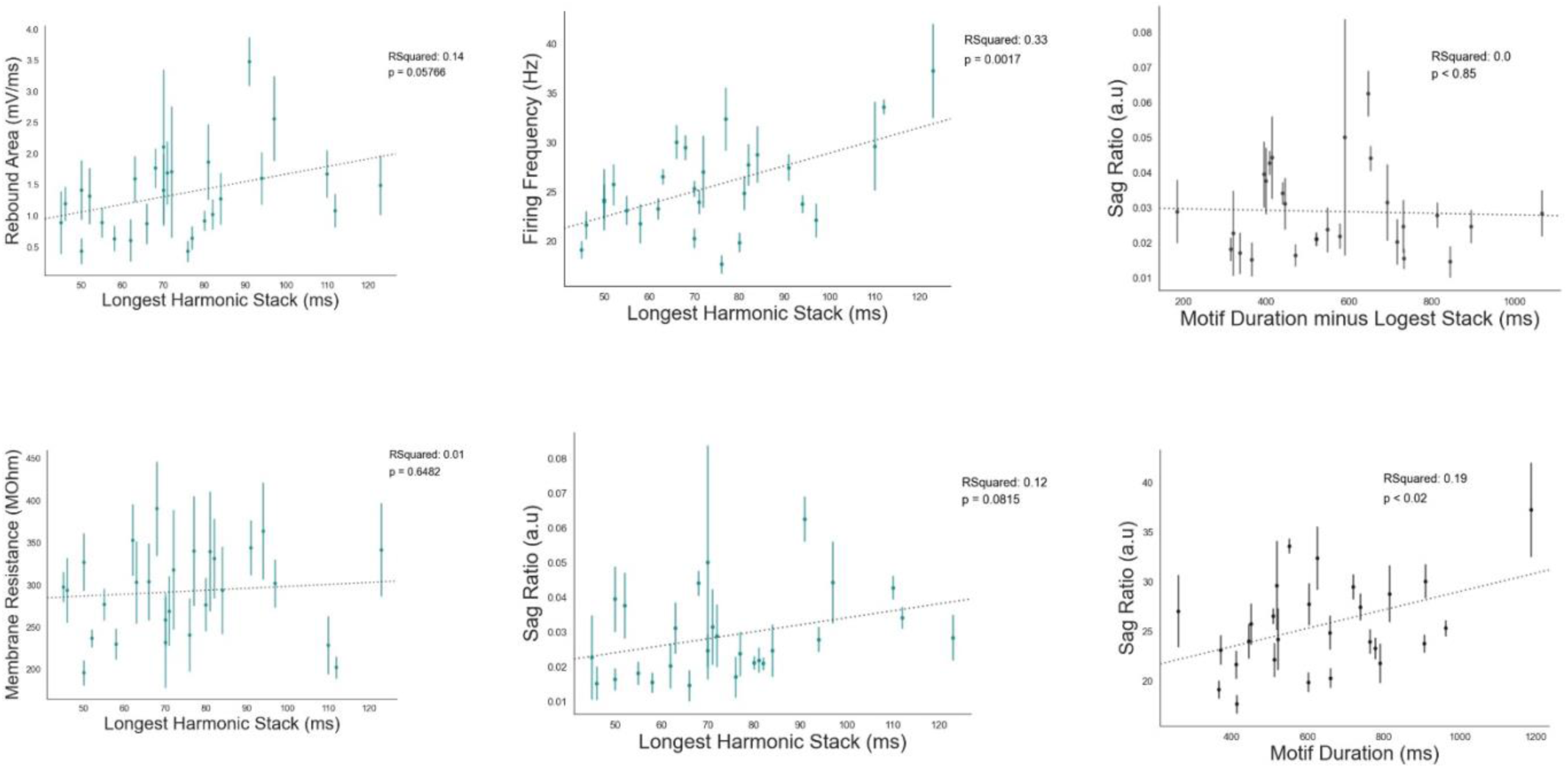
Relationships between Intrinsic properties and song features, excluding birds whose songs had long-duration (> 150 ms) longest harmonic stacks. Scatter plots of mean analyzed parameters for all HVC neurons for each bird, against features of song duration (error bars are standard error of the mean).

**Supplemental Figure 3.**
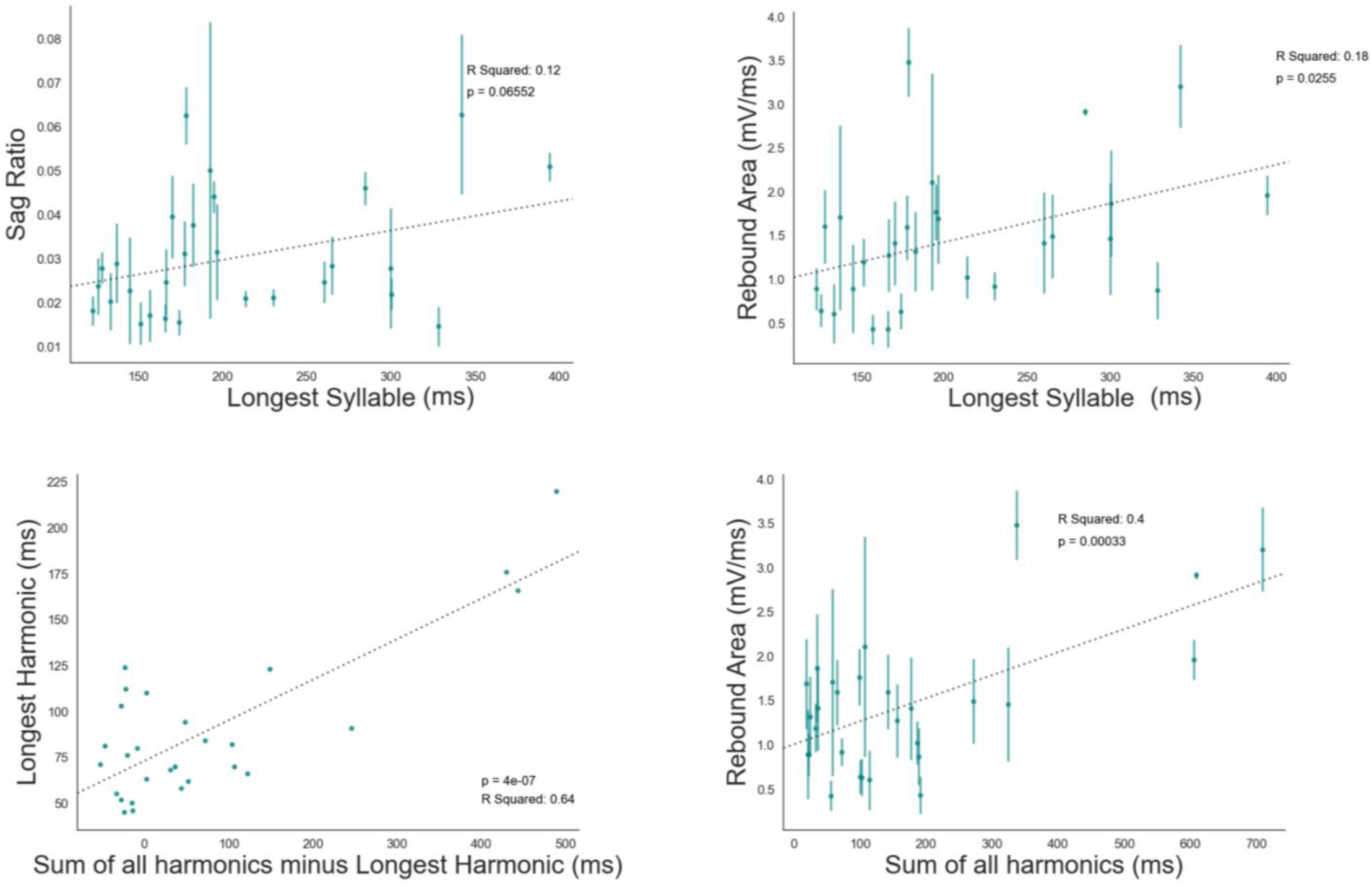
Additional correlations between intrinsic properties and temporal song structure. Scatter plots of mean analyzed parameters for all HVC_X_ for birds singing natural songs, against longest syllable and the sum of additional harmonic elements beyond the longest harmonic. Error bars represent standard error of the mean.

**Supplemental Figure 4.**
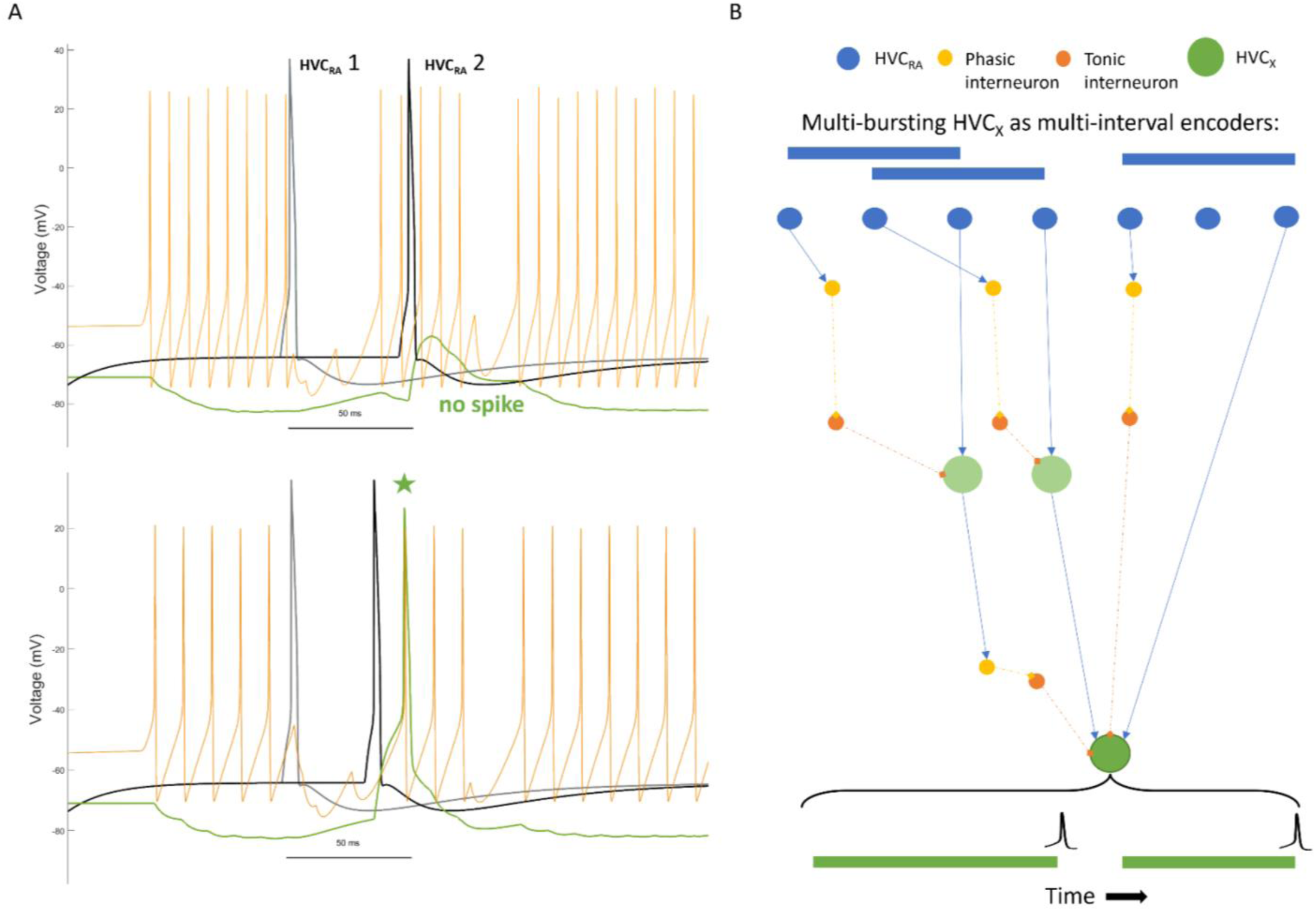
Single and multi-bursting model HVC_X_ neurons. A: Voltage traces from multiple neurons modeled and wired as described in Figure 5, illustrating the time dependence for the sequence sensitivity of the network model. An interneuron (orange trace) inhibits an HVC_X_ (green trace). One HVC_RA_ (first grey spike) di-synaptically inhibits the orange interneuron while a second, later-bursting HVC_RA_ (later black spike) excites the green HVC_X_ neuron. The top panel shows the outcome where the second spike arrives too late, resulting in no spike in the HVC_X_. The bottom panel shows a well-timed second HVC_RA_ spike producing a spike in the HVC_X_ (green star). **B:** Model circuit diagram depicting nested intervals leading to one HVC_X_ neuron (bottom dark green circle) that bursts twice.

**Supplemental Figure 5.**
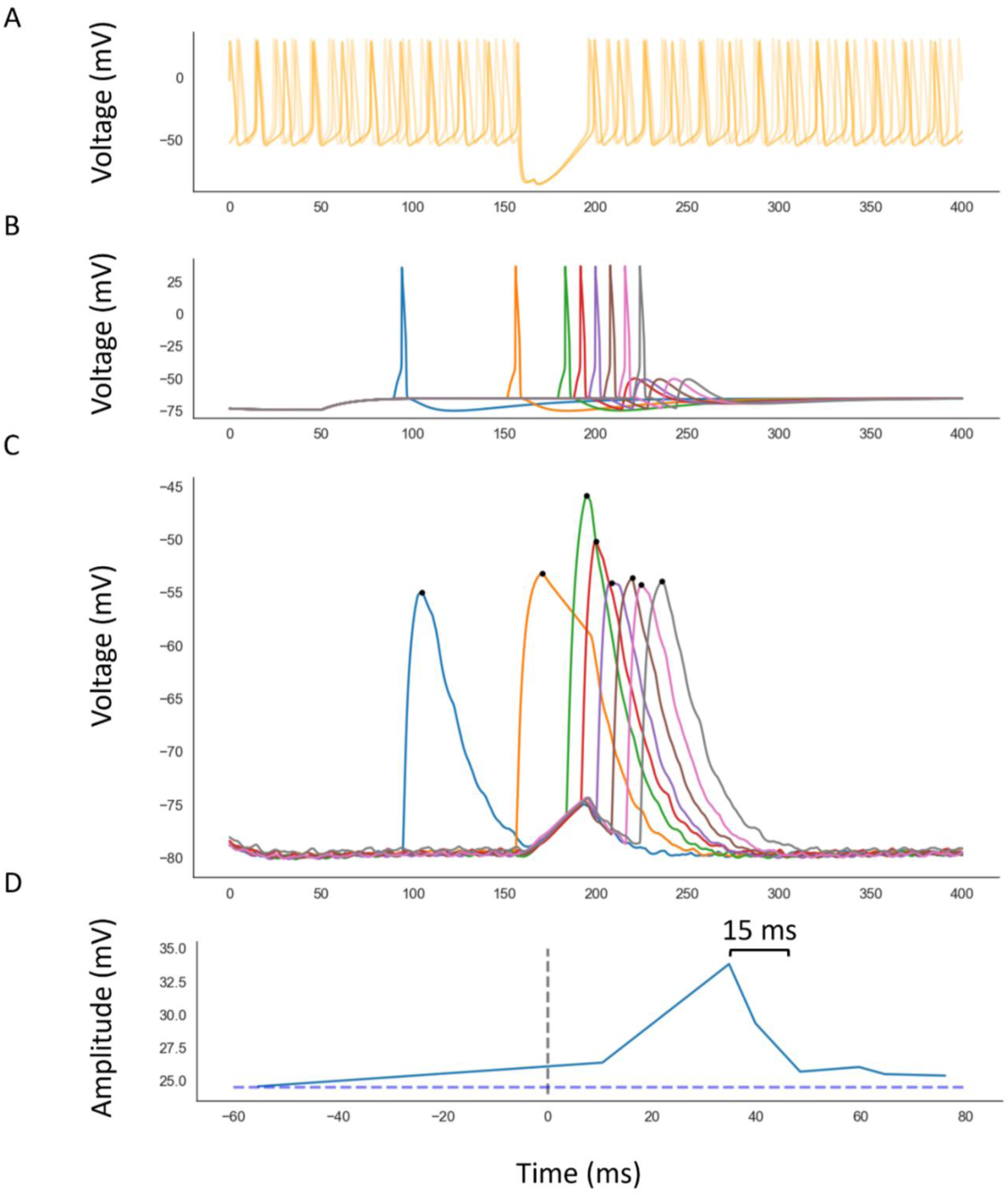
Time window encoded by HVC_X_. Hodgkin-Huxley model module with an HVC_X_ neuron with no voltage-gated sodium channels and varying timing of excitatory inputs. **A:** Overlayed traces from three interneurons synapsing onto one HVC_X_. **B:** Multiple HVC_RA_ voltage traces overlayed (each HVC_RA_ is depicted by a different color). **C:** The voltage traces of the same HVC_X_ arising from inputs from interneurons in A, and each individual HVC_RA_ in B. Peak voltages are shown by black dots. **D:** Peak amplitudes from C, and their relative timing from inhibition release (black dashed line). Baseline amplitude (blue dashed line) was taken from excitatory input that occurred before, and does not overlap with, release from inhibition.

**Supplemental Figure 6.**
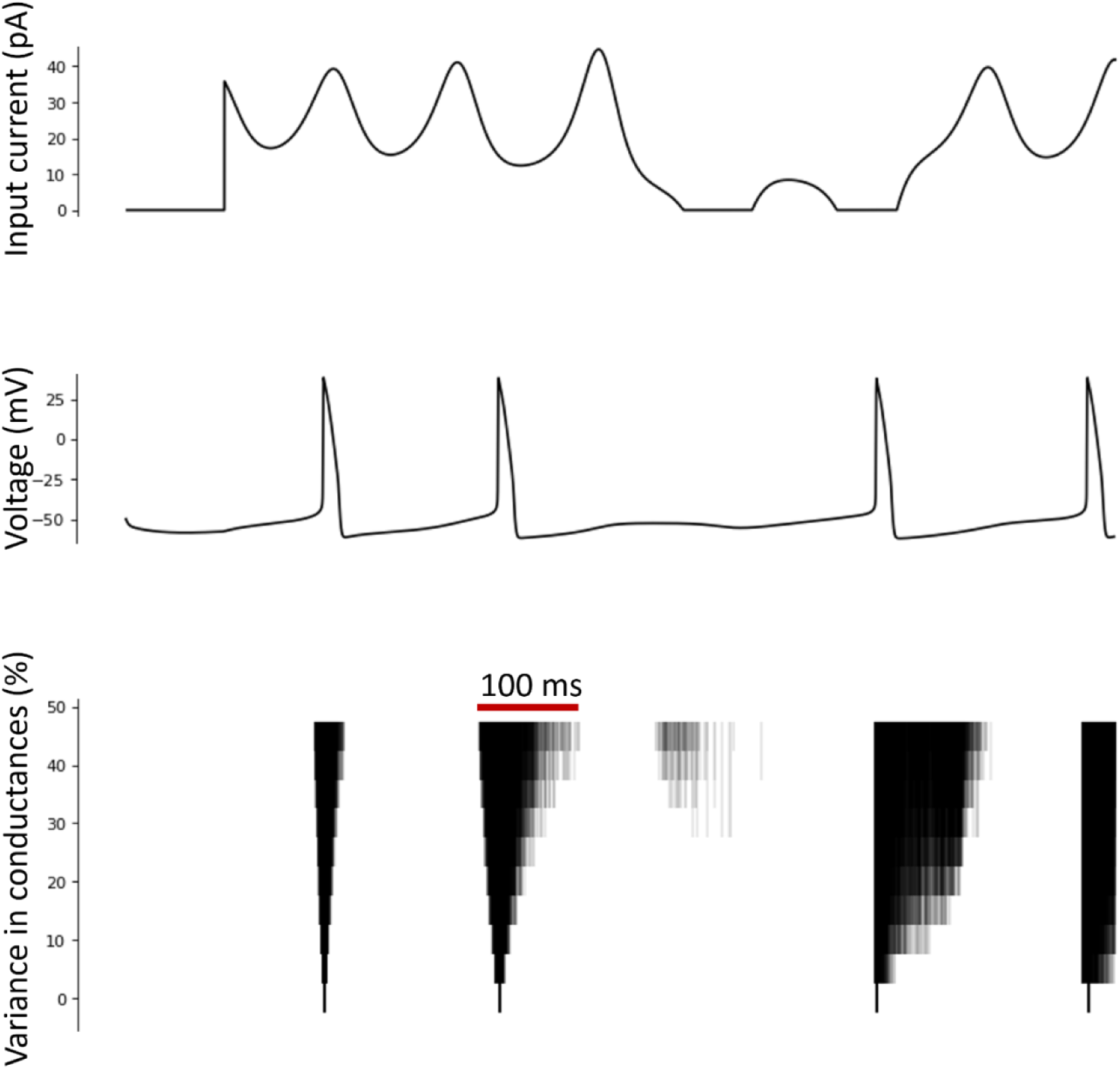
Intrinsic property homogeneity promotes spike time homogeneity in modeled neurons. Hodgkin-Huxley model neurons receiving identical inputs (top panel) and producing differently timed spike responses (one example model trace, middle panel). Adjusting percent variance among five modeled ionic conductances (gNa, gK, gH, gSK, and gCa-T) between 0 and 50% produced different ranges spike times (bottom panel). Each row in the bottom panel represents all spike times for 100 modeled neurons at a given range of IP variance.

**Supplemental Figure 7.**
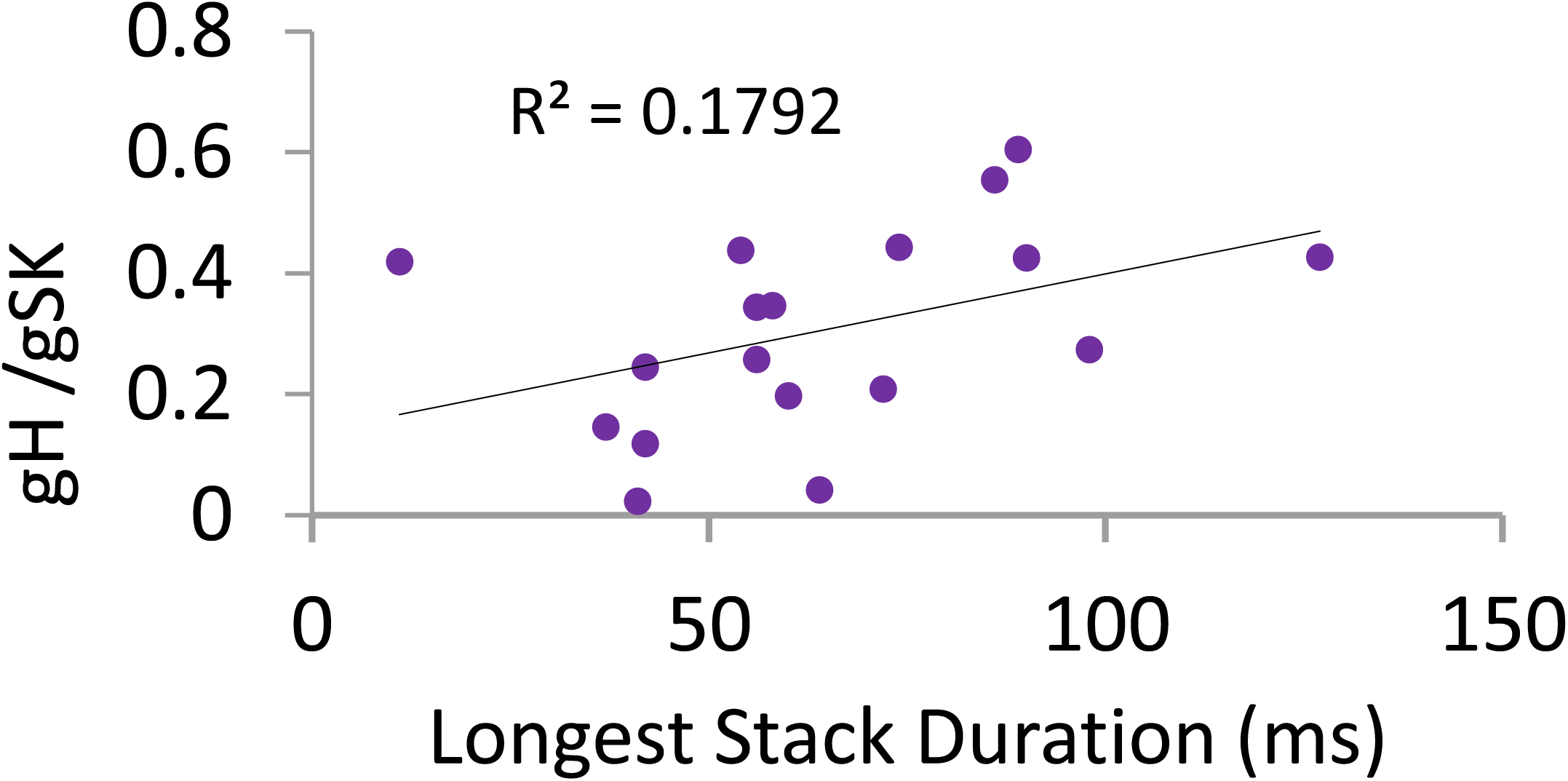
Relationships between modeled conductances and longest harmonic from Daou and Margoliash 2020. Scatter plots of mean gH divided by gSH for all HVC neurons for 18 birds.

